# The Mechanism of Histone Ubiquitylation by the ASB9-CUL5 Ubiquitin Ligase

**DOI:** 10.1101/2025.05.21.655409

**Authors:** Calvin P. Lin, Nathan H. Lee, Francis X. Alipranti, Harry Li, Elizabeth A. Komives

## Abstract

The E3 ligase substrate receptor ankyrin and SOCS box protein 9 (ASB9) was shown to bind over 10 different proteins including metabolic enzymes such as creatine kinase, filament proteins such as vimentin, and histones. In previous work, we characterized the ASB9-Cullin 5 E3 ligase (ASB9-CRL) ubiquitylation of creatine kinase and showed that ubiquitylation required the ring-between-ring ligase, ARIH2. Here we characterize the ASB9-CRL ubiquitylation of histones and show that histones H3 and H4 are polyubiquitylated by the ASB9-CRL whereas histones H2A and H2B are not. Many, but not all lysines in the histones are ubiquitylated suggesting some substrate specificity. Binding experiments show that the ligase-histone interaction is highly electrostatic and the neddylated ASB9-CRL binds with highest affinity. Only free histones are ubiquitylated. When the histones are in nucleosomes or in complex with the chaperone Asf1, they are not ubiquitylated. Only K48 and K63 polyubiquitin chains were observed, suggesting that the ubiquitylation probably drives protein degradation. The presence of ASB9 in specific cell types correlates with situations in which free histones H3 and H4 need to be degraded. In this work, we demonstrate that the ASB9-CRL is the ligase that facilitates degradation of histones H3 and H4. In addition, this work represents the first example of Cullin 5 mediated ubiquitylation that does not require a ring-between-ring “helper” ligase.

---

Ubiquitylation is a post-translational modification that regulates a variety of biological processes and is mediated through a multi-step enzymatic cascade (1). Most commonly, ubiquitin-conjugation mediates protein degradation required for regulating a multitude of biological processes such as gene transcription, the cell cycle, signaling cascades, viral immunity, and tumor suppression (2-4). The E1 ubiquitin (Ub) activating enzyme is first activated by Ub in an ATP-dependent manner, followed by Ub transfer to an E2 conjugating enzyme through a trans-thiolation reaction. E3 ubiquitin ligases then interact with Ub-conjugated E2s and substrate proteins to promote covalent Ub attachment to substrate ɛ-amino group of lysine residues by either a direct or indirect mechanism. In the direct mechanism, E3’s such as HECT (Homologous to the E6-AP Carboxyl Terminus) E3 ligases or RING-between-RING (RBR) ligases receive a Ub from an E2-Ub before ubiquitylating the substrate. In the more common indirect mechanism, E3s bind the E2-Ub and the Ub is transferred from the E2 to the substrate. In this mechanism, E3s such as Cullin-RING ligases (CRLs) act as a scaffold to allow ubiquitin conjugation from E2s to substrates. The human genome is comprised of two E1s, 37 E2s, and over 600 E3s to ubiquitylate a wide variety of proteins (5). The diversity of E3s is needed to recognize and ubiquitylate the diversity of cellular proteins (6).

The Cullin-RING ligase complex family, the largest group of E3 ligases, generally consists of similar key components including a substrate receptor, adaptor proteins, a Cullin, and a RING-Box protein (RBX) which binds an E2 carrying the thioester-linked Ub (7). Unique combinations of adaptor protein and substrate receptors allow CRLs to recruit and target thousands of substrates. Adaptor proteins interact with the N-terminal region of Cullins and recruit specific classes of substrate receptors depending on the interaction motif they contain. For example, Cullin 5 (CUL5) has been shown to bind Elongin B and Elongin C (ELOB/C), which then bind different ankyrin and suppressor of cytokine signaling (SOCS)-box (ASB) substrate receptors containing a consensus SOCS-box binding sequence (4). A quantitative SILAC mass spectrometry study comprehensively identified proteins that bound to each of the 18 ASB proteins (8). We decided to focus on the structurally characterized ASB9 (9-11). The ASB9 pulldowns confirmed interaction with proteins that constitute a Cullin5-RING ligase such as TCEB2 (Elongin B), TCEB1 (Elongin C), and CUL5. Creatine kinase B (CKB), a verified substrate of the ASB9-ELOB/C-CUL5-RBX2 (ASB9-CRL), also immunoprecipitated with ASB9 (12–16) in high abundance. Surprisingly, abundant interacting proteins varied in structure and spanned a variety of cellular functions such as metabolism (CKB, ENO2), membrane trafficking (RAB1A), and chromatin formation (H2AFZ, H2BFD, H3.3A, H4/A, and H1F2). ASB9 is found in only three main organs, kidney, liver, and most abundantly testis (12). Previous work from our lab developed a structural model of the ASB9-CRL with CKB bound from structures of CKB-ASB9-ELOB/C and CUL5-RBX2 (10) and showed that ubiquitylation of CKB requires the ring-between-ring ligase, ARIH2 (13). Here, we focus on determining whether histones are a substrate for the ASB9-CRL and, if so, whether the mechanism of ubiquitylation is the same or different from that of CKB.

Cullin-mediated substrate ubiquitylation often requires CRLs to partner with a RING-between-RING (RBR) E3 ARIH ligase family member (14), to form cooperative E3-E3 super-assemblies. One model suggests that certain folded Cullin substrates require an additional ARIH E3 ligase because the E2∼Ub located at the Cullin C-terminus is unable to reach the substrate near the Cullin N-terminus, while an ARIH∼Ub E3 ligase is positioned closer to such substrates(15). Mechanistic studies have only characterized CRL5 substrates that require ARIH2, specifically CKB ubiquitylation by ASB9-CRL (16, 17) and APOBEC3G ubiquitylation by Vif-CBFβ-CRL (13, 16, 18).

Histone accumulation (19, 20) and histone scarcity (21) have been shown to lead to genome instability and cell death. Thus, histone levels are tightly regulated through transcriptional, posttranscriptional, and posttranslational mechanisms including ubiquitylation. Histone ubiquitylation can occur in either DNA-bound or free histones that are not DNA-bound. CUL4 ligases have been shown to mediate histone mono- and poly-ubiquitylation in nucleosomes at sites of DNA damage (22) or chromatin remodeling (23) to release histones from DNA and recruit repair/remodeling proteins. Excess histones that accumulate in the cytosol have been shown to be ubiquitylated and degraded by HEL1 and HEL2 (24). Cullin 5 has been shown to localize to both the cytosol (14, 25) and the nucleus at sites of DNA damage (26) and transcriptional stalling (27).

Here we present *in vitro* studies of histone ubiquitylation with reconstituted ASB9-CRL to determine which histones are ubiquitylated and in what form (nucleosomal or free histones). Specific histone lysines that were modified was determined by the method of Gygi (28). These results identify ASB9-CRL as a novel histone E3 ligase and presents a model in which ASB9-CRL poly-ubiquitylates free histones H3 and H4 for proteasomal degradation.

## Experimental Procedures

### Expression Vectors

Vectors pertaining to ASB9-CRL complex, ubiquitin, E1 activating enzyme, and E2 conjugating enzyme expression in *E. Coli* were obtained as previously described (10, 17). Vectors for expression of *Xenopus laevis* histones H2A, H2B, H3, and H4 in *E. Coli* were a gift from Shannon Lauberth. Site directed mutagenesis of the WT H4 vector was done using blunt-end cloning to generate the H4 V22C vector. pQTEV-ASF1A was a gift from Konrad Buessow (Addgene plasmid # 31591; http://n2t.net/addgene:31591; RRID:Addgene_31591) (29).t

### E1/E2/E3 expression

Expression of ASB9 and ubiquitin machinery in *Escherichia coli* was performed as previously described (10, 17) with some modifications. Co-expression cells containing plasmids for ASB9/ELOB/C or CUL5/RBX2/ELOB/C were made using sequential transformation. The vector containing ELOB/C (Cam^R^) was transformed into competent BL21(DE3) cells (Invitrogen) after which those cells were made competent again. ELOB/C in pACYC was selected for by chloramphenicol (CAM) resistance. The vector for expression of ASB9 (Kan^R^) was transformed into ELOB/C-containing BL21 cells and plated on a kanamycin (Kan)-Cam LB agar plate for later culture inoculation and glycerol stock production. Similarly, vectors containing RBX2 (Amp^R^) and CUL5 (Kan^R^) were sequentially transformed into competent cells containing the ELOB/C vector (Cam^R^) to generate *E. coli* cells for co-expression of CUL5/RBX2/ELOB/C from cell plates or glycerol stocks. *E. coli* expression vectors containing RBX1 (Amp^R^) and CUL1 (Kan^R^) were sequentially transformed into BL21(DE3) cells for co-expression from KAN-AMP LB agar plates or from glycerol stocks. *E. coli* expression vectors containing Ub (Kan^R^) and UBE1 (Amp^R^) were sequentially transformed into BL21(DE3) cells for co-expression from KAN-AMP LB agar plates or from glycerol stocks. Vectors for expression of UBE2D2, UBE2L3, UBE2F, Ub, and NEDD8 were transformed individually into BL21 cells and plated on KAN LB agar plates. Vectors for expression of NAE1/UBA3 and ARIH2 were transformed individually into BL21 cells and plated on AMP LB agar plates. Transformation of the NAE1/UBA3 vector into competent cells containing the NEDD8 vector was performed as mentioned above for coexpression of NAE1/UBA3 and NEDD8.

All non-histone proteins were expressed as follows. A 20 mL M9-ZN (1.5 × M9 salts, NZ-Amine media, 0.8% dextrose, 1 mM MgSO_4_, 0.2 mM CaCl_2_) overnight culture was inoculated with a single colony from the plate or from a glycerol stock. The 1 L M9-ZN media was inoculated with the entire 20 ml starter culture and grown to OD_600_ = 0.8 at 37 °C. After placing the cultures on ice for 15 min, protein expression was induced by addition of IPTG to a final concentration of 0.5 mM, and the cultures were transferred to an 18 °C incubator for 20 h. RBX1, RBX2, and ARIH2 were expressed in media containing 200 μM ZnCl_2_ added just prior to induction.

### Histone expression

Vectors containing *Xenopus laevis* H2A, H2B, H3, and H4 were individually transformed into BL21 Star^TM^(DE3)pLysS (Invitrogen) and plated on AMP-CAM LB agar plates. Single colonies were used to inoculate 20 mL LB cultures and grown overnight with shaking at 37 °C. The overnight cultures were then used to inoculate 1 L large cultures of 2xYT media and grown to OD_600_ = 1.0 at 37 °C. Protein expression was induced by addition of IPTG to a final concentration 0.2 mM, and the cultures were grown for another 3 hours. Cells were collected by centrifugation at 5500 x g for 25 min and the collected pellet was frozen at -80°C until use.

### Protein Purification of ASB9-CRL5 Complexes, Ubiquitin Machinery, and Asf1

Purification of ASB9-containing complexes; ASB9-ELOB/C, ASB9-ELOB/C-CUL5-RBX2, or ubiquitin machinery; UBE1, UBE2D2, UBE2L3, UBE2F, and ARIH2 was performed as previously described (10, 17) with some modifications. In short, cells were resuspended in a buffer supplemented with protease inhibitor cocktail (Sigma P2714) and 5 mM PMSF, sonicated, and clarified by centrifugation at 25,000 xg for 35 minutes. The clarified lysate was incubated with 2 ml HisPur™ Ni-NTA Resin (Thermo 88221) (pre-equilibrated in resuspension buffer) for 2 h at 4 °C with rocking. Ni-NTA beads were pelleted by centrifugation at 800 x g for 5 min. The supernatant was discarded, and the Ni-NTA beads were poured into a glass Econo-Column® Chromatography Column (Bio-Rad), where Ni-NTA beads were washed with 20 mL wash buffer (50 mM Tris-HCl pH 8.0, 400 mM NaCl, 30 mM imidazole pH 8.0, 2 mM β-mercaptoethanol, 5% glycerol) at 4 °C, and His-tagged protein was eluted from the Ni-NTA beads using 14 mL elution buffer (50 mM Tris-HCL pH 8.0, 250 mM NaCl, 250 mM imidazole pH 8.0, 2 mM β-mercaptoethanol, 5% glycerol) over 30 min at 4 °C. The eluate was dialyzed overnight (10 kDa cut-off) in dialysis buffer (20 mM Tris-HCl pH 8.0, 100 mM NaCl, 5% glycerol, 1 mM DTT). Samples were concentrated to 2 mL and purified using size-exclusion chromatography (SEC) over a Superdex 200 16×600 column (Cytiva) in dialysis buffer. Peak fractions were combined and concentrated with Vivaspin® 10kDa MWCO concentrator (Sartorius) for use in experiments. Asf1 was purified in the same manner as described for the components of ASB9-containing complexes except without glycerol. The Asf1-H3-H4 complex was made by incubating the three proteins in a 2:1:1 ratio and purified by size-exclusion on a Superdex^TM^ 200 Increase 10/300 column.

### Neddylation of CUL5

ASB9-ELOB/C-CUL5-RBX2 (ASB9-CRL) was neddylated during its purification as previously described with some modifications (10, 17). Pellets of ASB9-ELOB/C (AE) (2/3 L cell pellet), ELOB/C-CUL5-RBX2 (ECR) (0.5 L cell pellet), and the NAE1/UBA3∼NEDD8 (0.5 L cell pellet) were resuspended in 100 mL resuspension buffer with protease inhibitor cocktail (Sigma P2714), 5 mM PMSF, and 10 mM MgCl_2_ and 2 mM ATP at 4 °C. Lysates were purified with 1 mL Ni-NTA resin as described above and dialyzed overnight. The dialyzed sample was brought to 5 mM MgCl_2_ and 2 mM ATP (Fisher BP413) by addition 1 M MgCl_2_ and a freshly made 100 mM ATP solution in dialysis buffer. 2 mg of previously purified ∼4 mg/mL UBE2F stocks in in 20 mM Tris pH 8.0, 250 mM NaCl, 50% glycerol, 1 mM DTT (the E2 for CUL5 neddylation) was added to the dialyzed sample and incubated at 4 °C for 2 h. The ∼8 mL sample was concentrated to ∼4 mL. The neddylated CUL5 complex was purified from NAE1/UBA3, UBE2F, and excess NEDD8 on a Superdex^TM^ S200 16x600 column (Cytiva) in dialysis buffer using a 2-mL injection loop. The fractions were analyzed by SDS-PAGE and fractions containing pure NEDD8-ASB9-ELOB/C-CUL5-RBX2 complex were combined, concentrated, and used immediately.

### Histone Purification

Histone monomers were prepared using established protocols (30, 31). Histones were expressed in BL21(DE3) pLysS *E. coli* cells and purified from inclusion bodies by gel-filtration using a Sephacryl 200 column (GE Healthcare), followed by dialysis in MQ-H_2_O containing 5 mM β-mercaptoethanol and lyophilization. Histones were resuspended in 0.5 M Na Acetate pH 5.2, 7 M urea, 200 mM NaCl, and 5 mM β-mercaptoethanol and further purified by ion-exchange chromatography using a 5-mL Hitrap SP HP column (Cytiva) followed by dialysis in MQ-H_2_O containing 5 mM β-mercaptoethanol, lyophilization, and storage at -80 °C.

### Histone Octamer Assembly and Purification

Histone octamers were prepared using established protocols (31, 32). Briefly, lyophilized histones were individually dissolved using 20 mM Tris-HCl pH 7.5, 7 M guanidine HCl, 5 mM β-mercaptoethanol to 2 mg/mL as determined by UV absorbance (ε_280_=44700 M^-1^cm^-1^). The four histone proteins were combined in equimolar amounts and adjusted to a final octamer concentration of 1 mg/mL. The solution was dialyzed against 10 mM Tris-HCl pH 7.5, 2 M NaCl, 1 mM DTT and octamers were purified from tetramer and dimer species using size-exclusion chromatography on a Superdex^TM^ 200 Increase 10/300 column (Cytiva). Pure fractions, as determined by SDS-PAGE, were combined and stored at -80 °C. Histone tetramers (H3-H4)_2_ were prepared the same way as octamers by using only lyophilized H3 and H4.

### Reconstitution of mononucleosomes

Mononucleosomes were formed as previously described (30-32) with some modifications. A plasmid containing 12 repeats of the Widom 601 DNA sequence (33) separated by 30 bp linkers was used to transform DH5α cells and the DNA was purified via phenol-chloroform extraction. DNA was digested into 177 bp repeats using ScaI (New England Biolabs) and purified. DNA pellets were dissolved in 10 mM Tris-HCl pH 8.0, 0.1 mM EDTA and stored at -20 °C. The DNA was mixed with histone octamers in a 1.2:1 ratio in 2 M TEK buffer (10 mM Tris-HCl pH 7.5, 2 M KCl, 0.1 mM EDTA, 1 mM DTT) and dialyzed into 10 mM Tris-HCl pH 7.5, 10 mM KCl, 1 mM DTT at 4 °C. Nucleosomes were analyzed on a APAGE (2% acrylamide, 1% agarose) gel (30).

### Experimental Design and Statistical Rationale

Having obtained all of the proteins and substrates necessary for the study, the experimental design involved three components. 1) At least two biological replicates of each *in vitro* ubiquitylation assay was performed and analyzed by SDS PAGE and mass spectrometry. ImageJ was used to measure the remaining ubiquitin in order to quantify the amount of ubiquitylation. Results are reported with standard deviations of the average of at least two independent experiments. 2) Bands and/or high molecular weight species were excised and subjected to in gel trypsin digestion followed by mass spectrometry. Two experiments were performed and three time points were collected. Only those peptides identified in at least two independent experiments and at least two time points are reported. 3) Fluorescence anisotropy was used to determine the binding affinity of the histones for the ASB9-CRL. In these experiments, the histones were fluorescently labeled at a specific cysteine. At least two experiments were performed and the average and standard deviations are reported.

### Nucleosome ubiqiutylation

Reactions to assess histone ubiquitylation in mono-nucleosomes were performed for 3 h at 37 °C with 0.2 μM UBE1, 0.5 μM UBE2D2, 50 μM Ub, 2 μM mono-nucleosomes, 1 μM ASB9-CRL, or 1 μM ASB9-CRL and/or 1 μM ARIH2-UBE2L3. Quenched reactions were run on a 13% SDS-PAGE gel. Two biological replicates were analyzed. Reactions comparing ubiquitin activity of ASB9-ELOB/C-CUL5-RBX2 (ASB9-CRL), CUL5-RBX2, CUL1-RBX1 were also done in a similar manner except incubation was only for 30 min with octamers and tetramers.

### Assessing histone ubiquitylation specificity

Reactions to assess ubiquitylation of free histones were carried out in a buffer containing 20 mM Tris-HCl pH 8.0, 200 mM NaCl, 5 mM MgCl_2_, 1 mM DTT, and 2 mM ATP with 0.2 μM UBE1, 0.5 μM UBE2D2, and/or ARIH2+UBE2L3, 50 μM Ub, and 0.5 μM NEDD8-ASB9-CRL or 0.5 μM ASB9-CRL with CUL5(K724R) at 37 °C. Reactions were initiated by addition of an E2-E3 master mix for concentrations of 0.5 μM UBE2D2 and 0.5 μM ASB9-CRL. Reactions were subsequently quenched at various time points by adding 2X SDS-PAGE buffer and boiling at 90 °C for 10 min prior to separation on a 4-20% Mini-PROTEAN® TGX™ SDS-PAGE gel. The degree of ubiquitylation was measured by quantifying band intensity of all unmodified histones using ImageJ (https://imagej.net/nih-image/). The mean and standard deviation were calculated from three biological replicates from different protein preparations.

Control reactions to assess the specificity of histone ubiquitylation were performed for 30 min at 37 °C in the same buffer with 0.2 μM UBE1, 0.5 μM UBE2D2, and 50 μM Ub as controls with either 4 μM H2A, 4 μM H2B, 4 μM H3, 4 μM H4, 4 μM H2A/H2B dimers, or 2 μM histone octamers. Reactions were initiated by adding 0.5 μM NEDD8-ASB9-CRL. The presented gel is representative of 2 biological replicates. Two biological replicates were analyzed.

Reactions with the Asf1-H3-H4 complex were performed in 20 mM Tris-HCl pH 8.0, 300 mM NaCl, 5 mM MgCl_2_, 1 mM DTT, and 2 mM ATP.

### Binding assays using fluorescence anisotropy

Fluorescently-labeled histone monomers were prepared as follows. Single lyophilized histone monomers of H3 C110 or H4 (V22C) were resuspended at a concentration of ∼2 mg/mL in unfolding buffer (20 mM Tris-HCl pH 7.5, 7 M guanidine HCl, 25 mM NaCl, 1 mM DTT). Oregon Green™ 488 Maleimide (OG488, Invitrogen^TM^ O6034), was prepared as a 100 mM stock in dimethylformamide and added to the histone suspension to achieve one molar equivalent and allowed to incubate overnight at 4°C. To remove remaining free OG488, samples were injected onto a Superdex^TM^ 200 Increase 10/300 column (GE Life Sciences) equilibrated in unfolding buffer using a 500 μL sample loop at 4 °C. Fractions were analyzed by SDS-PAGE and fractions containing pure labeled histones were pooled dialyzed, lyophilized, and stored at -80 °C.

Labeled histones were dissolved in unfolding buffer at ∼1 mg/mL and dialyzed overnight into buffers of various salt concentrations (20 mM Tris-HCl, 10 mM NaCl or 150 mM NaCl, 5% glycerol, 1 mM DTT) just prior to fluorescence anisotropy experiments. Protein concentration and labeling efficiency was assessed by measuring the absorbance at 280 nm and 495 nm on a Nanodrop spectrophotometer.

Black 96-well plates (Greiner Bio 675076) were first prepared by incubating 20 μL 5.5 mg/mL BSA in each well for 20 minutes at 25 °C. OG488-labeled histones (5 nM) were mixed with various concentrations of ASB9-CRL or NEDD8-ASB9-CRL complexes in triplicate in the coated wells and incubated at 25 °C for 1.5-2.5 h. Fluorescence anisotropy was measured at 25 °C on a SpectraMax iD5 Multi-Mode Microplate Reader (Molecular Devices) using an excitation wavelength of 485 nm, an emission wavelength of 535 nm, and an integration time of 100 ms. Anisotropy was calculated using r = [I_(V,V)_ − GI_(V,H)_]/[I_(V,V)_ − 2GI_(V,H)_], where r is anisotropy, I_(V,V)_ is the fluorescence intensity in the parallel direction, I_(V,H)_ is the fluorescence intensity in the perpendicular direction, and G is the grating factor used for calculating the anisotropy is 1.0, as previously determined for the instrument measurement pathway.

Fluorescence anisotropy data for the histones were fit to the following equation to determine the equilibrium binding affinity:

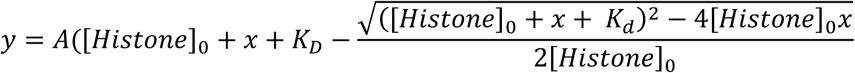

where y is the change in fluorescence anisotropy, A is the maximum change in fluorescence anisotropy, x is the varying concentration of ASB9-CRL5 complex, [Histone]_0_ is the concentration of histone (5 nM), and K_D_ is the equilibrium binding affinity (33). A values were determined for each individual set of triplicates due to slight variability in the anisotropy plateau values.

At least two biological replicates for each condition were conducted on separate days with different histone and ASB9-CRL5 complex preparations. For each assay, data were fit to the aforementioned equation to determine the K_D_. The reported K_D_ values are the mean and SEM of the K_D_ values determined for each of the three replicate experiments. Normalized data from all 3 sets of triplicates were plotted.

### Ubiquitylation analysis by mass spectrometry

The aforementioned ubiquitylation reactions were carried out for 2, 5, and 10 min, quenched by addition of gel-loading buffer, and electrophoresed on a 13% Tris-glycine SDS-PAGE gel and stained with Coomassie Brilliant Blue G-250 (Thermo Scientific™ 20279). Gel bands were excised and cut into small pieces and washed according to standard procedures (34). In-gel proteins were acetylated with 5 μL acetic anhydride and brought to pH ∼7 with ∼150 uL 1 M ammonium bicarbonate followed by 1 h incubation at 37 °C. Then, cysteines were reduced and alkylated with 1.36 mg/mL chloroacetamide in 100 mM ammonium bicarbonate, 8 mM TCEP, washed and immediately digested with trypsin for 30 min at 4 °C, then 37 °C overnight. The supernatant containing tryptic peptides was collected (34). Samples were dried in a speed-vac and stored at -20 °C until analysis.

Trypsin-digested peptides were analyzed by ultra high pressure liquid chromatography (UPLC) coupled with tandem mass spectrometry (LC-MS/MS) using nano-spray ionization on an Orbitrap fusion Lumos hybrid mass spectrometer (Thermo) interfaced with nano-scale reversed-phase UPLC (Thermo Dionex UltiMate™ 3000 RSLC nano System) using a 25 cm, 75-micron ID glass capillary packed with 1.7-µm C18 (130) BEHTM beads (Waters corporation). Peptides were eluted from the C18 column into the mass spectrometer using a linear gradient (5–80%) of Acetonitrile (ACN) at a flow rate of 375 nl/min for 1.5 h. The buffers used to create the ACN gradient were: Buffer A (98% H_2_O, 2% ACN, 0.1% formic acid) and Buffer B (100% ACN, 0.1% formic acid). Data dependent acquisition mode was used for data collection and the mass spectrometer parameters were as follows; the MS1 survey scan using the orbitrap detector (mass range (m/z): 400-1500 using quadrupole isolation, 60000 resolution setting, spray voltage of 2200 V, ion transfer tube temperature of 275 °C, AGC target of 400000, and maximum injection time of 50 ms). Data dependent scans (top speed for most intense ions, with charge state set to only include +2-+5 ions, and 5 second exclusion time, while selecting ions with minimal intensities of 50000 at in which the collision event was carried out in the high energy collision cell (HCD Collision Energy of 30%), and the fragment masses were analyzed in the ion trap mass analyzer (with ion trap scan rate of turbo, first mass m/z was 100, AGC Target 5000 and maximum injection time of 35ms). Protein identification was carried out using Peaks Studio X (Bioinformatics solutions Inc.). Search parameters were as follows: Precursor Mass Error Tolerance: 10.00ppm, Fragment Mass Error Tolerance: 0.02Da, Max Missed Cleavage: 2, Max Variable PTM per Peptide: 2. Carbamidomethylation (+57.02) was the only fixed modification and variable modifications included: acetic anhydrate (+42.01), oxidation (M) (+15.99), ubiquitin (+114.04), and ubiquitination (+383.23).

Ubiquitin modifications were observed through trypsin products “GG” or “LRGG” left on ubiquitylated lysines, with masses of 114.04 or 383.23 Da respectively. A cutoff score of (-10logP) > 14 (p-value < 0.040) was used to validate PTMs. Ubiquitylation of lysines was monitored over 2, 5, and 10 min ubiquitin reactions. Only those modifications that were observed in at least two of the three timepoints are reported.

## Results

### Nucleosome core particles are not the preferred substrate for ASB9-CRL

Because all core histones were highly enriched in an ASB9 pull-down (8), initial ubiquitylation assays were performed with mononucleosomes as substrates. Mononucleosomes were assembled using previously established methods using a 177 bp DNA sequence comprised of a high-affinity 147 bp Widom sequence with a 30 bp linker and histone octamers. Ubiquitylation reactions containing mono-nucleosomes were incubated with ASB9-CRL at 37°C but no ubiquitylation was observed even after 3 h (**Figure 1**). Addition of ARIH2 and its obligate E2, UBE2L3, to reactions with ASB9-CRL and mononucleosomes still did not elicit any histone ubiquitylation.

**Figure 1.**
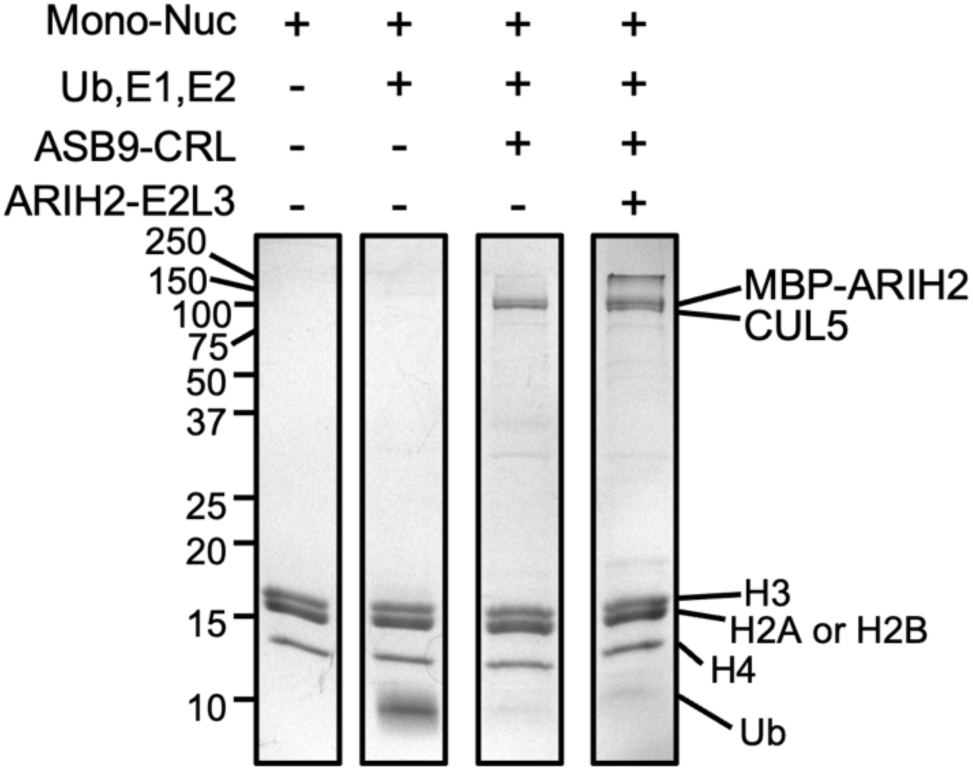
ASB9-CRL does not ubiquitylate histones in mononucleosomes. ASB9-CRL ubiquitylation assays using nucleosomes with or without an additional E3 ligase, ARIH2 required for ubiquitylation of another ASB9 substrate, CKB. Representative image of n = 2 independent experiments.

### Histones H3 and H4 are poly-ubiquitylated by the ASB9 Cullin5 E3 ligase

We next attempted to ubiquitylate extranucleosomal histones in the absence of DNA. Reactions were performed with histone octamers against ASB9-CRL with and without a NEDD8 post-translational modification (NEDD8ylation) of Cullin5 which has been shown to enhance Cullin-mediated substrate ubiquitylation (35-37). Assays with NEDD8-ASB9-CRL showed mono- and di-ubiquitylation of histones H3 and H4 by 2 minutes as identified by mass spectrometry (**Figure 2A, Table S1**). Polyubiquitylation of histones H3 and H4 was also very rapid (**Figure 2A, Table S2**). To analyze the effect of neddylation on ASB9-CRL ubiquitylation, ubiquitylation was performed with a ASB9-CRL reconstituted with a Cullin5 K724R mutant that cannot be neddylated, showing that NEDD8 promotes histone ubiquitylation (**Figure 2B**). Previous studies showed dissociation of histone octamers into H2A/H2B dimers and H3 and H4 monomers (38) which would be expected in our experimental conditions, so it is possible that the ASB9-CRL only ubiquitylates monomeric histones. To test this hypothesis, ubiquitylation assays with NEDD8-ASB9-CRL were performed on individual histone monomers, H2A/H2B dimers, and histone octamers. Again, the most poly-ubiquitylated species were produced from histones H3 and H4 (**Figure 2C**). Full ubiquitylation was confirmed by the complete disappearance of the unmodified monomeric H3 and H4, whereas monomeric histones H2A and/or H2B remained mostly unmodified. Moreover, there appeared to be almost no ubiquitylation of the H2A/H2B dimer. These results indicated that ASB9-CRL histone ubiquitylation is specific to H3 and H4. Finally, when the histone chaperone, Asf1 was in complex with histones H3 and H4, no histone ubiquitylation was observed. This result further substantiated that free, extranucleosomal H3 and H4 were the preferred substrates of ASB9-CRL (**Figure S1**).

**Figure 2.**
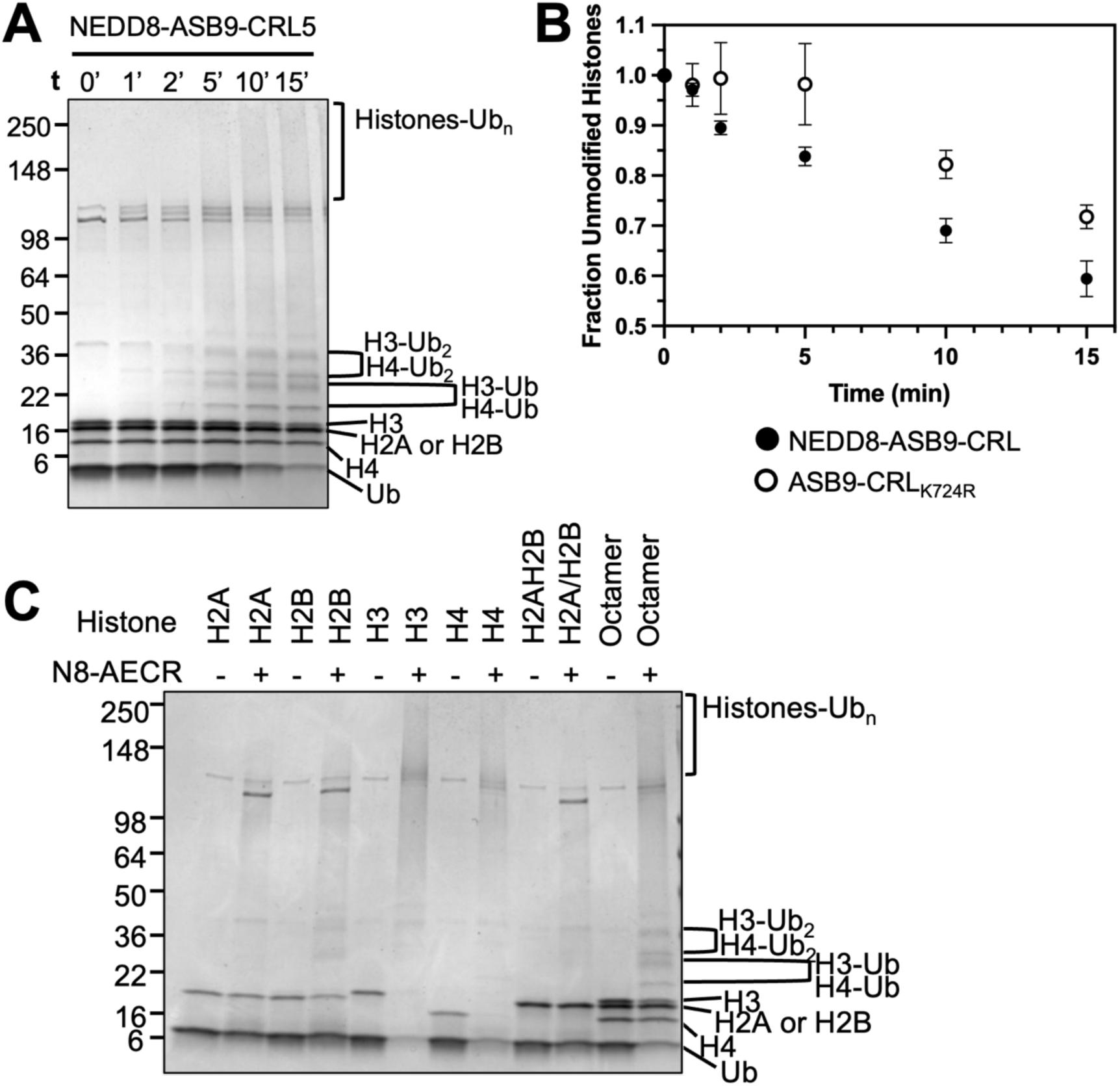
*In vitro* ubiquitylation assays of histone octamers shows specificity towards Histones H3 and H4. (**A**) Time course of histone octamer ubiquitylation by NEDD8-ASB9-CRL. Covalently ubiquitin-modified histones appeared as higher weight mono-ubiquitylated bands, di-ubiquitylated bands, and poly-ubiquitylated high-molecular weight smears. (**B**) Histone octamer ubiquitylation assay with NEDD8-ASB9-CRL and an unneddylated ASB9-CRL using a K724R CUL5 mutant showed that neddylation increased histone ubiquitylation. ImageJ was used to quantify band intensity of all remaining unmodified histones from biological triplicates using separate protein preparations. The fraction of unmodified histones approached 0.5 because the gel band of histones H2A/H2B, which ran together, did not appear to decrease over time. (**C**) 30 min ubiquitylation reactions of individual histones, H2A/H2B dimer, and histone octamers. H3 and H4, including those in the octamer sample shows a preference towards H3 and H4. Reactions with H2A and H2B, as monomers or dimers, showed minimal ubiquitylation. Images are representative of n = 3 biological replicates.

Reactions which did not include ASB9 or Elongins B/C had some activity, similar to reactions with CUL1-RBX1 as quantified by measuring band intensity of unmodified histones in reactions containing H3 and H4 against a control without ligase components (**Figure S2A-B**), where using the full ASB9-CRL had notably enhanced H3 and H4 ubiquitylation. Taken together, these results strongly suggest that ASB9-CRL is a specific E3 ligase for histones H3 and H4.

### Identification of poly-ubiquitylated histone species

To determine which histone lysines were rapidly ubiquitylated, ubiquitylation reactions of histone octamers were performed with NEDD8-ASB9-CRL at 2, 5, and 10 min. Reactions were quenched with SDS PAGE loading buffer and fractionated on 13% SDS-PAGE gels followed by in-gel lysine acetylation, trypsin digestion, and LC-MS/MS analysis to identify the ubiquitylated histone species. The mono- and di-ubiquitylated species labeled in **Figure 2A** were confirmed, and histone species in high molecular weight gel bands were identified (**Figure 3**). Ubiquitin modifications were detected through peptides with a modification of 114.04 or 383.23 Da which corresponds to tryptic ‘GG’ or ‘LRGG’ residues on ubiquitylated lysines. Unmodified lysines were identified by acetylation as indicated by a modification of 42.01 Da (**Figure 3**). Mass spectrometry identified histone ubiquitylation almost exclusively in histones H3 and H4, further solidifying our conclusion that ubiquitylation is specific to histone H3 and H4. Data analysis across all three time points showed high reproducibility for ubiquitylated lysines (**Table S2**). Mass spectrometry also identified ubiquitylated lysines for Cullin 5 indicating auto-ubiquitylation as previously observed (7, 39). While ubiquitin chains made by UBE2D2 are suggested to not have a linkage specificity (40), mass spectrometry indicated exclusively K48 and K63 linkages. It is important to note that we do not know that all of the lysines that were ubiquitylated were polyubiquitylated because the mass spectrometry data gives the aggregate result. It is possible that only one or two of the lysines were polyubiquitylated and that was sufficient to result in a high molecular weight histone species while other lysines may have been only mono or di-ubiquitylated. The mass spectrometry data is insufficient to ascertain which lysines were actually polyubiquitylated.

**Figure 3.**
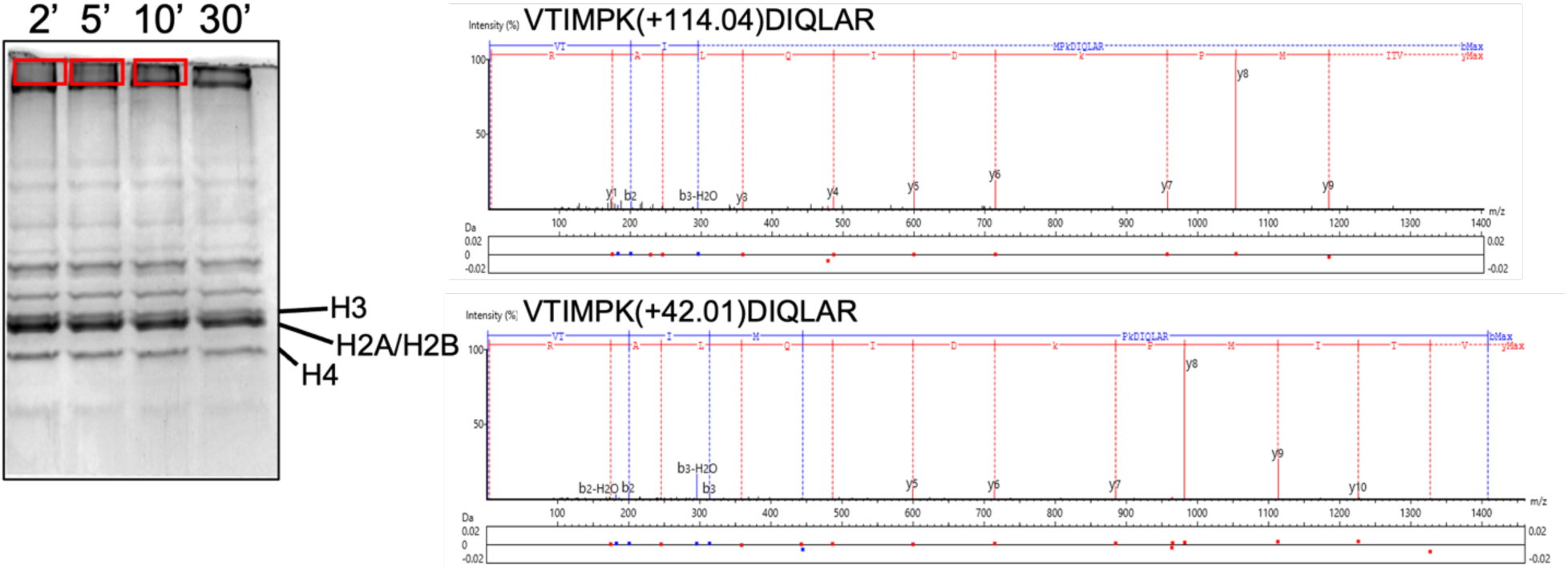
Mass spectrometry identifies specific ubiquitylated lysines. *In vitro* ubiquitylation PTM analysis workflow. The same 2, 5, and 10 min ubiquitylation reactions in Figure 2 of histone octamers with NEDD8-ASB9-CUL5-ELOB/C-RBX2 were run on a 13% SDS-PAGE gel for 45 minutes and stained with Coomassie Blue. Individual bands of polyubiquitylated species from the gel (marked by boxes) were excised, acetylated, alkylated, and trypsin digested before analysis using LC-MS/MS. Two example MS2 spectra of ubiquitylated and acetylated (unmodified) H3 peptide 117-128 are shown, illustrating identification of modification. Here, the m/z differences between the y6 and y7 ions reveal m/z changes relative to unmodified peptides used to identify post-translational modification.

Mapping of the ubiquitylated lysines onto the primary sequence and structures of H3 and H4 predicted by Alphafold (**Figure 4 A, B**) showed that ubiquitylated lysines were present in both the disordered histone tails (H3K18, H3K23, and H4K31), and the core histone fold (H3K56, H3K79, H3K122, H4K31, H4K77, and H4K91). Importantly, ubiquitylation was not observed in some lysines that were covered in the mass spectrometry data (H3K9, H3K14, H3K36, H3K37, H3K64, H4K20, H4K79), indicating that the ASB9-CRL has some apparent substrate specificity for which lysines it modifies. These results confirm that the ASB9-CRL ubiquitylates free histones H3 and H4 at specific sites. The in-gel digest mass spectrometry results reveal that while a small amount of histones H3 and H4 remain mono- or di-ubiquitylated, the majority of histones H3 and H4 are rapidly poly-ubiquitylated within 2 minutes at specific lysines by ASB9-CRL.

**Figure 4.**
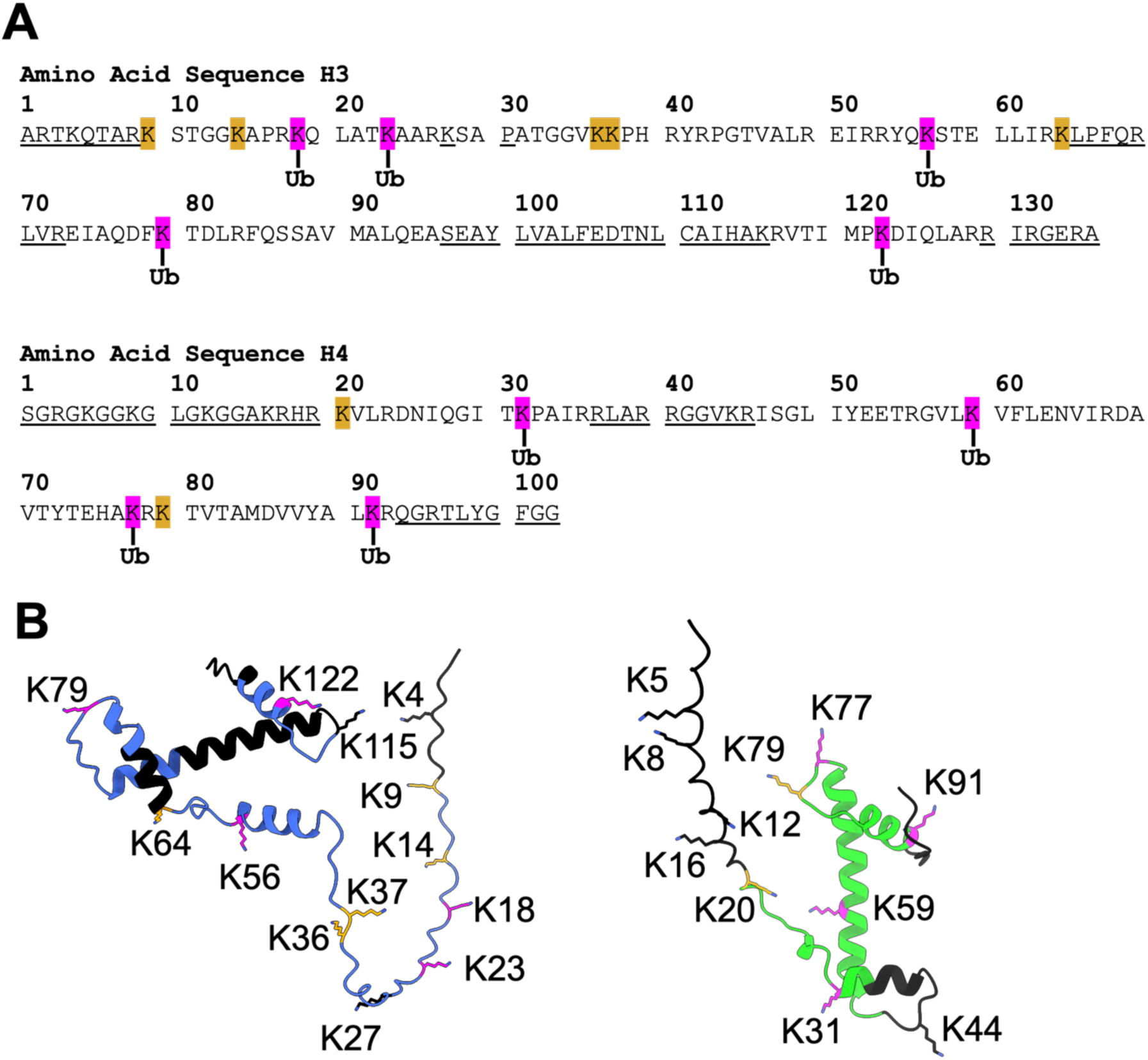
Histones H3 and H4 ubiquitylation pattern. (**A**) Primary amino acid sequence of histones *Xenopus laevis* H3 and H4. Identified lysines modified with ubiquitin are shown in magenta. Unmodified lysines that were covered by MS data were shown in orange, while sequences that were not covered by mass spectrometry were underlined. (**B**) Structures of histones H3 (blue) and H4 (green) were predicted by Alphafold and the identified lysines were mapped onto the structure (modified with ubiquitin in magenta, unmodified lysines in orange, and regions unidentified by MS in black).

### Quantitative determination of H3 and H4 binding affinity to ASB9-CRL ligases

To measure equilibrium binding affinities of NEDD8-ASB9-CRL and ASB9-CRL to histones H3 and H4, fluorescence anisotropy assays were performed. The single cysteine in histone H3, C110, and installed cysteine in histone H4, V22C, were fluorescently labeled with the cysteine-reactive Oregon Green™ 488 Maleimide, purified, and a constant amount of each labeled histone was mixed with increasing amounts of NEDD8-ASB9-CRL, ASB9-CRL, or CUL5-RBX2. Preliminary experiments revealed a strong electrostatic dependence on the binding with affinities, as decreasing the salt concentration 10 mM NaCl significantly enhanced ASB9-CRL binding of H3 and H4 compared to experiments performed at 150 mM NaCl which did not produce quantifiable binding affinities (41). ASB9-CRL bound to H3 and H4 with K_D_ of 187 ± 38 nM and 79.7 ± 6.8 nM respectively and neddylated ASB9-CRL showed an approximate 2-fold enhancement of binding affinity with K_D_ of 78.2 ± 8.7 nM and 48.4 ± 9.7 nM respectively (**Figures 5A, B**). Binding affinities with only the substrate receptor and adaptor protein(s) subcomplex ASB9-ELOB/C, were too weak and could not be accurately measured, which was unlike previous studies showing ASB9-CRL substrate CKB binding to ASB9 with sub-nanomolar affinity (42). Intriguingly, CUL5-RBX2 alone elicited binding to histones H3 and H4 albeit weaker than the full neddylated and unneddylated ASB9-CRL ligases (**Figure 5A, B**).

**Figure 5.**
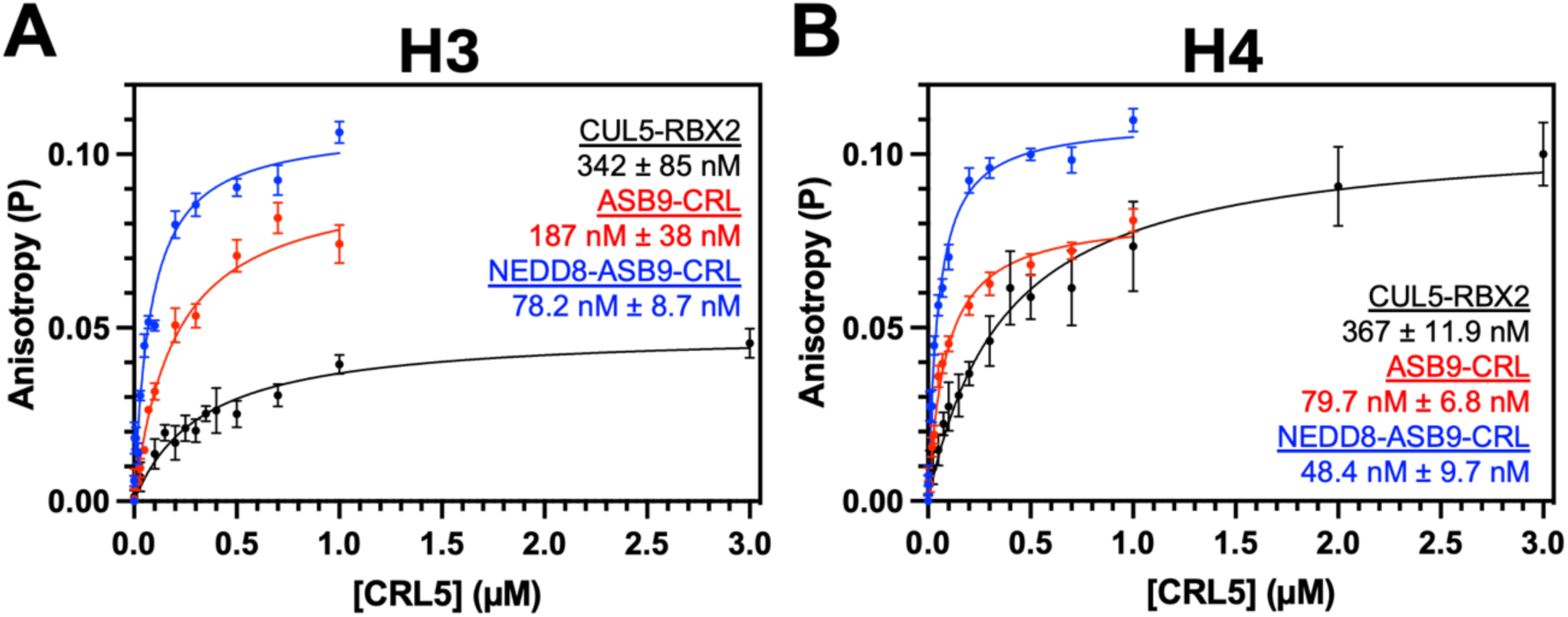
Fluorescence Anisotropy of histones H3 and H4 with CRL. Fluorescence anisotropy with Oregon Green 488 labeled histones (**A**) H3 and (**B**) H4V22C showed nM affinity against CUL5-RBX2, ASB9-CRL, and NEDD8-ASB9-CRL at 10 mM NaCl. Slight binding enhancements for both histones were observed upon neddylation of CUL5. Data represent the mean and SEM of at least two biological replicates and the reported K_D_ values are the mean and SEM of K_D_ calculated from each individual experiment.

### ARIH2 is not required for histone ubiquitylation

Previous studies showed that the RBR ligase, ARIH2 was essential for ASB9-CRL ubiquitylation of CKB (16, 17). To test if ARIH2 participated in NEDD8-ASB9-CRL ubiquitylation of histones, ubiquitylation assays were performed with ARIH2-UBE2L3 added to the NEDD8-ASB9-CRL. No differences were observed in the amount of ubiquitylation as compared to reactions with only NEDD8-ASB9-CRL (**Figure 6**), showing that histone ubiquitylation is not ARIH2 dependent.

**Figure 6.**
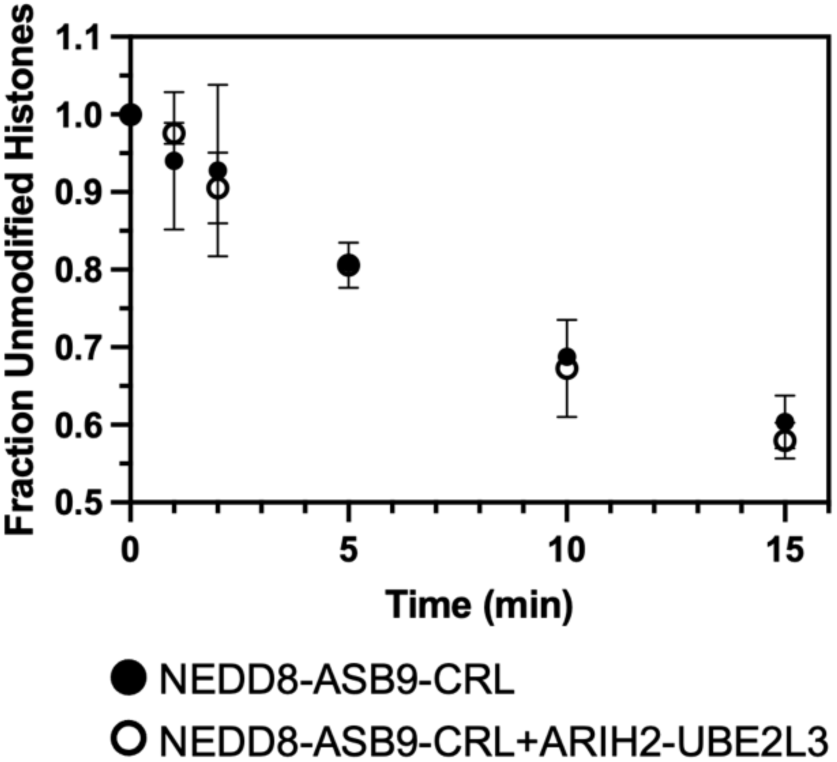
ARIH2 is not required for ubiquitylation of histones. The RBR ligase ARIH2 along with E2 UBEL23 have been shown to be required for ubiquitylation of certain CRL5 substrates. Ubiquitylation reactions of octamers with NEDD8-ASB9-CRL ± ARIH2-UBE2L3 did not affect ubiquitylation. ImageJ was used to quantify band intensity of all remaining unmodified histones from two biological replicates using separate protein preparations.

## Discussion

Our results show that ASB9-CRL polyubiquitylates histones H3 and H4 but not histones H2A and H2B. Given that the other *bona fide* substrate of the ASB9-CRL is the metabolic enzyme, creatine kinase B, this finding reveals that the ASB9-CRL can ubiquitylate structurally diverse targets. This contrasts with other substrate receptors, such as the βTrCP receptor, which specifically binds conserved phosphorylated degron sequences (43).

We showed that ASB9-CRL-mediated polyubiquitylation of histones does not require ARIH2, in contrast to the requirement of ARIH2 for ubiquitylation of CKB. In fact, all Cullin 5-containing CRL ubiquitylation reactions so far have been shown to require ARIH2. It is thought that well-folded substrates are too far from the Ub∼E2s in CRLs and this is why they require an additional ring-between-ring ligase (13, 15). Extranucleosomal, non-chaperone bound histones exist in an extended α-helical form under near-physiological salt conditions (200 mM NaCl) (44-46), and may be disordered enough that they don’t require ARIH2 for ubiquitylation. Structuring the H3 and H4 by Asf1 chaperone binding or nucleosome complexation completely inhibits ASB9-CRL ubiquitylation.

*In vitro* assays offered insights into how ASB9-CRL recognizes and targets H3 and H4 for ubiquitylation. The binding affinity of histones H3 and H4 for the ASB9-CRL was in the nanomolar range at 10 mM NaCl and weakened to micromolar at physiological salt concentrations indicative of an electrostatically driven protein-protein interaction. Neddylation of the Cullin 5 was shown to enhance binding affinity to histones H3 and H4. The substrate receptor ASB9 was insufficient for even micromolar affinity binding to histones H3 and H4, whereas CKB bound so tightly to ASB9 that no dissociation was observed in SPR experiments (42). Surprisingly, CUL5-RBX2, even without a substrate receptor, bound directly to histones H3 and H4, although with four-fold lower affinity than NEDD8-ASB9-CRL and two-fold lower affinity to ASB9-CRL. It is therefore evident that ASB9-ELOB/C works synergistically with CUL5-RBX2 as a complex to facilitate histone ubiquitylation, potentially with CUL5-RBX2 being involved in direct binding of histones.

Mass spectrometry analysis of the high molecular weight H3 and H4 species contained K48 and K63 ubiquitin linkages showing that histones H3 and H4 were rapidly polyubiquitylated (within 2 min) by the ASB9-CRL. Since the functional consequence of K48 and K63 linked polyubiquitylation is proteasomal degradation, it is likely that ASB9-CRL polyubiquitylation of histones H3 and H4 targets them for proteasomal degradation (47, 48).

Ordinarily, histones are found stabilized in nucleosomes. All eukaryotes have multiple genes encoding histones and have the potential to generate excess histones (49), resulting in deleterious effects including cell death (20). During UV damage, histones H3 and H4 are ubiquitylated (50). CUL5 has been shown to translocate into the nucleus to sites of DNA damage (27) and pol II stalling (26). Another CRL, CUL4-DDB-ROC1, has been shown to ubiquitylate nucleosomal histones, as part of the DNA damage response. UV-irradiation has shown that histones H3 and H4 are mono-ubiquitylated by CUL4, which causes them to be ejected from nucleosomes, and facilitates subsequent recruitment of the pyrimidine dimer repair protein XPC-RAD23B (22). It is possible that mono-ubiquitylation destabilizes the nucleosomes releasing the histones for ASB9-CRL polyubiquitylation.

DNA damage can also cause an accumulation of excess cytoplasmic histones (24, 51, 52). Previously, yeast have been shown to combat cytoplasmic histones through proteasomal degradation mediated by RING ligases, HEL1 and HEL2, and E2 Ubc4 (homologous to human UBE2D2) (53) and Ubc5 (homologous to UBE2D1). Thus, cytoplasmic ubiquitylation of free histones serves as a potential function of ASB9-CRL since CUL5 is present in the cytoplasm and its E2, UBE2D2, was shown to be required for ubiquitylation of free histones in yeast. Moreover, the polyubiquitylated histones K48 and K63 linkages observed by mass spectrometry suggests a proteasomal-degradation function (47, 48). In yeast, various chaperone proteins interact with free histones to eliminate non-nucleosomal histone-DNA interactions, particularly upon DNA damage (54). We tested whether the ASB9-CRL could degrade chaperone (Asf1)-bound histones H3 and H4 but it could not. Since the ASB9-CRL is found in both the nucleus and the cytoplasm, it could play a role in removing free histones in both locations during DNA damage.

The first report of H3 ubiquitylation showed polyubiquitylation of H3 in elongating spermatids of rat testes (23). Although a role was not assigned to polyubiquitylation of H3, it was suggested that degradation may facilitate chromatin remodeling where DNA is further compacted in the sperm’s head by protamines (55). In fact, testis is the organ showing the highest level of ASB9 expression (12) and expression of ASB9 increases over the maturation of spermatogonial stem cells into spermatids (56, 57). Low expression of ASB9 was observed in patients with abnormal spermatogenesis (56). Thus, ASB9-mediated degradation is essential for normal sperm maturation.

In conclusion, our results show, for the first time, that ASB9-CRL specifically polyubiquitylates histones H3 and H4 revealing a new function for the ASB9-CRL. Ubiquitylation of free histones has been shown to be important for degradation of histones in normal spermatogenesis, the tissue where ASB9 expression is highest. Although all other Cul5 substrates known so far require the additional ring-between-ring ligase, ARIH2, our results show that the ubiquitylation of histones H3 and H4 by ASB9-CRL does not require ARIH2.

## Abbreviations

SOCS: Supressor of Cytokine Signaling
ASB9: Ankyrin and SOCS box protein 9
CRL: Cullin Ring Ligase (here Cullin is Cullin 5)
H2A: Histone H2A
H2B: Histone H2B
H3: histone H3
H4: histone H4
CKB: creatine kinase brain-type
ELOB/C: elongin B/elongin C
RBRLs: RING between RING E3 ligases
Ub: ubiquitin

## Data Availability

All mass spectrometry data is available at MassIVE (massive.ucsd.edu) with the data set MSV000097954 and password a.

## Supplemental data

This article contains supplemental data

## Acknowledgments

We thank Majid Ghassemian of UCSD’s BPMSF for assistance with mass spectrometry and Ryan J. Lumpkin for mentorship and insightful discussion regarding CRL’s. This work was supported by NSF grant MCB 1817774. C.P.L. was supported by the Molecular Biophysics Training Grant, NIH Grant T32 GM00832.

## Notes

https://massive.ucsd.edu/ProteoSAFe/private-dataset.jsp?task=247635203c184831b63315d8031dff34

### Competing Interest Statement

The authors have declared no competing interest.

