## Supplementary Figures for "The Mechanism of Histone Ubiquitylation by the ASB9-CUL5 Ubiquitin Ligase"

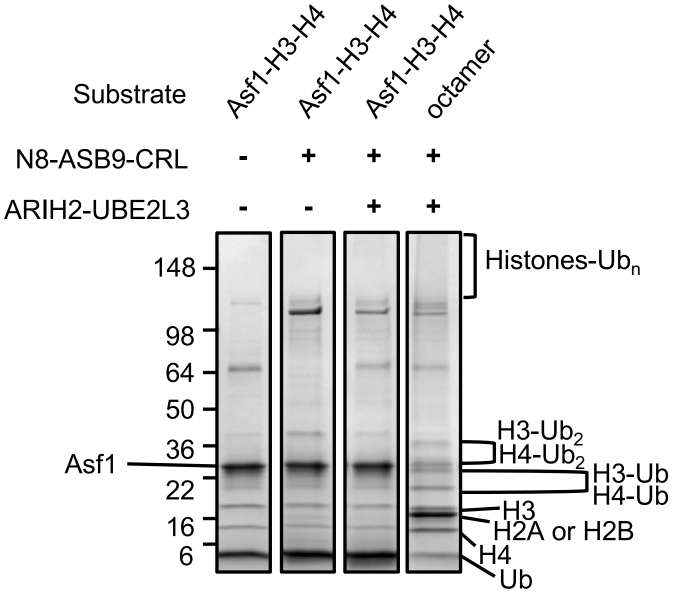


**Supplementary Figure 1** – Ubiquitylation reactions with H3-H4-Asf1. Extranucleosomal histones H3 and H4 in complex with chaperone Asf1 were not ubiquitylated by N8-ASB9-CRL, while inclusion ARIH2-UBE2L3 similarly had no effect in contrast with octamer ubiquitylation. Gel Image is representative of n=2 biological replicates.

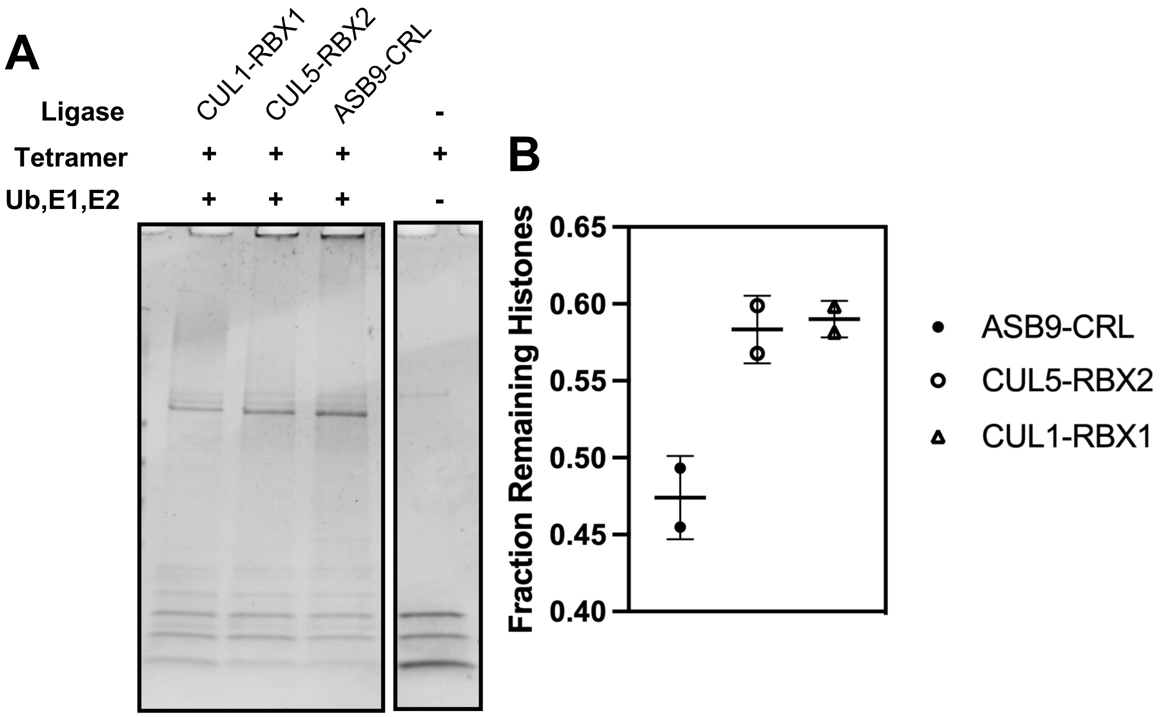


**Supplementary Figure 2** – Ubiquitylation of H3 and H4 with various Cullin ligases. (**A**) Cul5-Rbx2 and Cul1-Rbx1 yields minimal ubiquitylation of H3 and H4, while inclusion of ASB9-EloB/C enhances ubiquitylation. (**B**) Histone ubiquitylation quantified against a control reaction (rightmost lane in **Figure S2A**) containing no ubiquitin machinery. Data is representative of at least two independent, biological replicates.
